# Multiple evolutionary routes to cytoskeletal arborization revealed by the rhizarian amoeba *Filoreta ramosa*

**DOI:** 10.1101/2025.10.24.684464

**Authors:** Sarah L. Guest, Scott C. Dawson

**Affiliations:** University of Massachusetts Amherst, Department of Biology, 611 N. Pleasant St., Amherst MA 01003; University of California, Davis; Department of Microbiology and Molecular Genetics One Shields Avenue, Davis, CA 95616

## Abstract

The eukaryotic cytoskeleton generates remarkable diversity in cellular architecture despite being built from deeply conserved actin and tubulin polymers. Diversification of cytoskeletal regulators, motors, and filament-organizing proteins produces highly varied cellular morphologies across eukaryotes, yet certain higher-order architectures repeatedly emerge in distantly related lineages. One notable example is cytoskeletal arborization, which occurs not only in metazoan neurons but also in amoeboid lineages distributed throughout the eukaryotic tree. Whether these similar branched architectures arise through conserved cytoskeletal organization, independent reuse of shared molecular systems, or convergence driven by common physical constraints remains unresolved. Here, we investigate the rhizarian amoeba *Filoreta ramosa*, which forms a multinucleate reticulated network through branching and anastomosis. Using live imaging, immunofluorescence, morphometric analyses, and cytoskeletal drugs, we define how actin and microtubule systems organize branch formation, intracellular transport, and large-scale network architecture. Actin-rich protrusions initiate exploratory branchlets that become selectively stabilized through microtubule incorporation. Longitudinal microtubule arrays reinforce mature branches and support rapid bidirectional organelle transport, while branch nodes function as distributed sites of microtubule nucleation. These cytoskeletal features parallel key mechanisms underlying neuronal arborization, including actin-driven exploration, microtubule-dependent branch stabilization, and transport systems that scale with increasingly extended cytoskeletal networks. However, unlike neurons, *Filoreta* develops a decentralized reticulated network through repeated anastomosis and distributed microtubule organization, demonstrating that similar arborized morphologies can emerge through distinct architectural strategies. Our findings indicate that arborization can arise through multiple evolutionary adaptations to common cellular constraints. Shared cytoskeletal mechanisms repeatedly support branching architectures, but distinct topologies and modes of cellular organization demonstrate that evolution can reach arborization through different routes. Similar cytoskeletal networks may repeatedly emerge in diverse lineages when cells face the challenges of exploration, stabilization, and transport across increasingly larger scales. *Filoreta* therefore provides an experimentally tractable model for investigating how conserved cytoskeletal systems generate diverse arborized cellular architectures across eukaryotic evolution.

## INTRODUCTION

The eukaryotic cytoskeleton exhibits remarkable diversity in organization, dynamics, and function across microbial and multicellular lineages. Although actin and tubulin polymers are deeply conserved throughout eukaryotes, the architectures they generate vary dramatically between cell types and organisms. This structural and functional diversity emerges not simply from the polymers themselves, but from extensive diversification of cytoskeletal binding proteins, motors, nucleators, crosslinkers, and regulatory factors that control filament assembly, stability, organization, and force generation (1–5). Cytoskeletal evolution is therefore thought to proceed through lineage-specific gene loss, duplication, neofunctionalization, and rewiring of regulatory systems. Conserved cellular functions such as intracellular transport or protrusion formation can consequently emerge through distinct molecular implementations, including convergently evolved MyTH4-FERM myosins and expanded kinesin repertoires in land plants following dynein loss (6–8). Despite this extensive cytoskeletal protein diversification, certain higher-order cytoskeletal architectures repeatedly emerge across deeply divergent eukaryotic lineages.

One striking example is arborized or branched cytoskeletal morphology or arborization. Although neurons are among the best-characterized branched cell types, branching and anastomosing pseudopods occur across multiple eukaryotic clades, including Amoebozoa, Stramenopiles, Rhizaria, and Opisthokonts (9–11). Similar branched architectures also arise during epithelial and tracheal morphogenesis, where coordinated actin remodeling and microtubule stabilization generate extended cytoskeletal networks (12,13). The repeated emergence of arborized morphologies despite extensive cytoskeletal diversification raises a fundamental evolutionary question: do similar branching architectures emerge through conserved cytoskeletal organization, recurrent reuse of shared molecular systems, or convergence driven by common physical constraints?

Large cells face fundamental organizational challenges involving surface area expansion, intracellular transport, spatial coordination, and environmental interaction. Arborization provides a scalable solution to these constraints by increasing surface area while maintaining connectivity across extended cytoskeletal networks. This organization also creates new logistical demands, as increasingly elaborate arbors require efficient intracellular transport and structural stabilization over long distances. Modern studies of branching morphogenesis demonstrate that these demands are frequently solved through coordinated interactions between actin-driven protrusion and microtubule-based stabilization (14,15). In many systems, direct coupling between actin assembly and microtubule dynamics guides branch positioning, stabilization, and persistence (16,17). If arborization represents a recurring cytoskeletal adaptation for increasing cellular scales, then similar coordination between exploratory actin protrusions and microtubule stabilization should emerge repeatedly across divergent lineages. Similar arborized architectures may therefore arise through multiple evolutionary routes, even when they are assembled from different molecular repertoires.

Amoebae provide a powerful comparative system for investigating these questions because their morphologies emerge directly from cytoskeletal organization. Unlike cells constrained by rigid walls or multicellular tissue architecture, amoeboid cell shape arises dynamically through cytoskeletal forces acting on the plasma membrane. Amoebae therefore provide an opportunity to directly examine how cytoskeletal systems generate complex cellular architectures. The Rhizaria are a major eukaryotic clade comprising diverse free-living and symbiotic protists that are abundant in marine and freshwater ecosystems and play important roles in global carbon cycling and microbial food webs (18,19). Many rhizarians possess highly elaborate cytoskeletal systems that generate thin filose pseudopods, reticulated networks, axopodia, and mineralized cellular structures, making the group a rich source of morphological and cytoskeletal diversity (20,21). Despite this diversity, the cytoskeletal mechanisms underlying rhizarian morphology remain poorly understood due to the limited availability of experimentally tractable model systems.

Reticulose rhizarians are particularly informative because they combine extreme cellular scale with highly branched morphologies and rapid intracellular transport. Classical studies of *Reticulomyxa filosa* demonstrated rapid bidirectional organelle transport along longitudinal microtubule arrays within a highly dynamic reticulated network (22–24). These studies established that large syncytial amoebae could support extensive microtubule-based transport systems but did not resolve how arborized architectures themselves are generated, stabilized, and reorganized. As a result, the cytoskeletal mechanisms that generate and organize large arborized cellular networks remain poorly understood.

Here, we investigate the rhizarian amoeba *Filoreta ramosa*, a recently isolated species whose life cycle includes a multinucleate syncytial stage that develops an extensive reticulated network through branching and anastomosis. Using live imaging, immunofluorescence, morphometric analyses, and pharmacological perturbations, we characterize how actin and microtubule systems organize branch formation, intracellular transport, and large-scale network architecture. Actin-rich protrusions initiate exploratory branchlets that become selectively stabilized through microtubule incorporation, while longitudinal microtubule arrays reinforce mature branches and support rapid bidirectional organelle transport. We further demonstrate that branch nodes function as distributed sites of microtubule nucleation, revealing a decentralized mode of cytoskeletal organization distributed throughout the syncytial network. These findings directly connect cytoskeletal organization to arborized morphogenesis in a rhizarian lineage and provide experimental support for the broader notion that arborized cytoskeletal architectures can emerge independently in diverse eukaryotic lineages through shared basic mechanisms despite differences in network topology. Together, these findings suggest that similar arborized morphologies can evolve through multiple adaptations to common cellular constraints, with conserved ancestral cytoskeletal mechanisms repeatedly modified to generate diverse forms of branched cellular organization across eukaryotes.

## RESULTS

### Dynamic, reticulated branched networks define the development of the *Filoreta* syncytium

In initial observations of *Filoreta* in culture, we observed an extensive reticulated network of dynamic branches and loops that alter morphology in response to environmental cues or stimuli. To define the precise nature of the branching development of this expansive arborized syncytium, we performed time-lapse imaging that captured key events and cellular behaviors in *Filoreta* branched syncytial development throughout the life cycle. In a growing syncytium, numerous peripheral pseudopodia extend as transient “branchlets” (**Figure 1A**, **1G**) that extend outward. Branchlets protrude as wide, flat, ruffled lamellar pseudopodia or thin, branched filopodia. The branchlets also formed loops by a process termed anastomosis (**Figure 1B**). Both filose and lamellar pseudopodial branchlets frequently collapsed and folded in on themselves, but branchlets that anastomosed were often more stable and persisted during outgrowth. These branchlet intersections became “nodes” in a reticulate syncytium (**Figure 1G**). Branchlets at nodes became the thicker stabilized “branches” of the network as the syncytium grew outwards (**Figure 1G**). Stable branches did not retract or fold back in on themselves and remained part of the network topology. Serial repetition of this branching, extension, and fusion process yielded a reticulate morphology.

**Figure 1:**
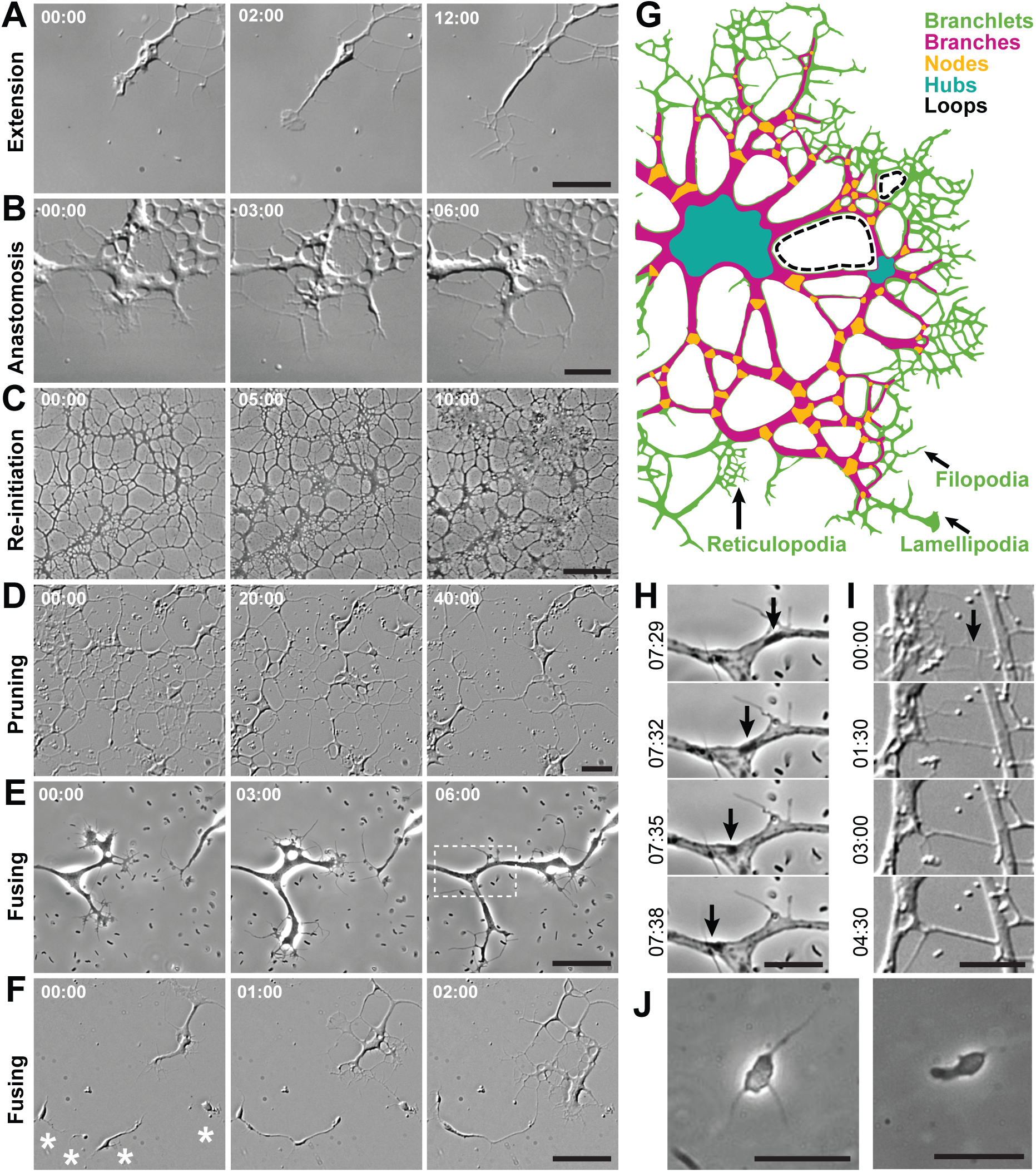
Dynamic, reticulated branched networks define the development of the *Filoreta* syncytium. Live syncytia were imaged during different developmental stages. (**A**) Pseudopodial branchlets extend as they crawl outwards, constantly growing and retracting as lamellipodia and filopodia. Left panel = ruffled pseudopod, center = lamellipodia, right = branched filopodia. Scale = 10μm. (**B**) Branch self-fusion (asterisk) is known as anastomosis, which generates the looped structure of the syncytial network. Scale = 20μm. (**C**) The syncytial network is dynamic and responds to the environment, re-initiating pseudopodia from simplified nodes following addition of nutrients to the media. Scale = 100μm. (**D**) The syncytium responds to stress by pruning back its branches and retracting into hubs. Syncytia were stressed by long-term incubation without nutrient-rich media and exposed to bright light. Scale = 20μm. (**E**) Syncytial neighbors recognize and fuse to each other. The inset rectangle in the right panel was used in (**H**). Scale = 10μm. (**F**) Individual amoebae fuse to each other and are recruited by a nearby syncytium. Scale = 20μm. (**G**) A colored diagram showing the morphology and associated terminology used to describe syncytial networks. Branchlets (green) = transient pseudopodial extensions that may be filopodia, lamellipodia or reticulopodia (bottom, arrows). Branches (magenta) = non-transient thicker lengths that make up the network. Nodes (orange) = intersecting points of two or more major branches that occur via bifurcation or anastomosis. Hubs (blue) = broad regions of cell body with five or more major branches intersecting. Loops (black) = closed circles occurring through anastomosis that may either be complete (filled in by branches, larger dotted region), or incomplete (partially bordered by branches or only by branchlets, smaller dotted region). (**H**) A montage from inset in (**E**) with frame intervals of 3 seconds show an organelle traversing a branch node (arrows). Scale = 5 µm. (**I**) A montage with frame intervals of 90 seconds showing branchlet to branch transition and thickening (arrow). Scale = 20µm. (**J**) An individual amoeba forming filopodia (left) and a small lamellipodia-like pseudopod (right). Scale = 10μm.

We also examined *Filoreta* network growth in response to varying nutrient levels. When grown in enriched media, the reticulated network structure developed continuously across the entire culture surface area. To test the dynamic response of stable branches, we treated established networks with 0.02% yeast extract and tryptone (YET), which increases nutrient concentration by twofold. Within five minutes after treatment, the network altered its branch topology by reinitiating new branchlets and lateral pseudopod extensions (**Figure 1C**). Newly initiated branchlets thickened and became part of the interior network topology, increasing the density of branch coverage on the culture surface (**Figure 1I**). In contrast to increasing branch density, syncytia under starvation or stress decreased their branch density. “Pruning” is a phenomenon of branch simplification and loss described as a part of the maturing process in dendritic arbors (25). During pruning, *Filoreta* networks simplified branch topology by retraction (**Figure 1D**). These observations indicate *Filoreta* continuously detects changes in its environment and rapidly alters its network in response.

Adding to network plasticity, syncytia also grew toward each other and fused (**Figure 1E, Video S1**). Neighboring syncytia fused together to form a larger syncytium, after which reorganization of the shared cytoskeletal components and organelles was observed. Organelles moved bidirectionally in branches throughout the network and caused the membrane to bulge as they traversed narrow branches (**Figure 1H**). Anastomosed filopodia within the same syncytia or between neighboring syncytia transitioned from “branchlets” to “branches” by a distinctive thickening of the adjoined filopodia. When grown in cultures with five times the normal nutrient concentration (0.05% YET), individual amoebae were present and readily fused to the network (**Figure 1F**). Individual amoebae were approximately 10 µm in diameter and crawled using filopodia and lamellar ruffles (**Figure 1J**). When in close proximity to each other, or to a nearby syncytium, their pseudopods elongated until they contacted the neighboring cell, immediately fused and were incorporated into the network. These observations suggested that network elaboration depends on dynamic transitions between transient exploratory protrusions and stabilized branches, prompting us to directly examine the cytoskeletal organization underlying branch development.

### Syncytia have distinct cytoskeletal architectures in pseudopodial branchlets and branches

To determine the underlying cytoskeletal architecture driving the dynamic morphology in *Filoreta*, we fixed and stained the actin and microtubule arrays in syncytia at different stages of development, using phalloidin and tubulin antibody. Phalloidin staining revealed dense actin filaments and patches in filopodial branchlets, minor and major branches (**Figure 2A, 2B, 2C**). These pseudopodial projections were particularly actin-enriched in branch fusion sites with filopodia, reticulopodia (**Figure 2A**), and lamellar pseudopodia (**Figure 3A**). Branches comprising the interior loops of the network contained longitudinal microtubules (**Figure 2C**). At the transition area between branches and branchlets, longitudinally oriented microtubules were partially incorporated into a fraction of branchlets (**Figure 2B, 3C**). The major branch “hubs” that become sites of macrocyst formation contain microtubule ends and are often highlighted by actin-based ruffled structures similar to lamellipodia observed at the hub boundaries (**Figure 2D**).

**Figure 2:**
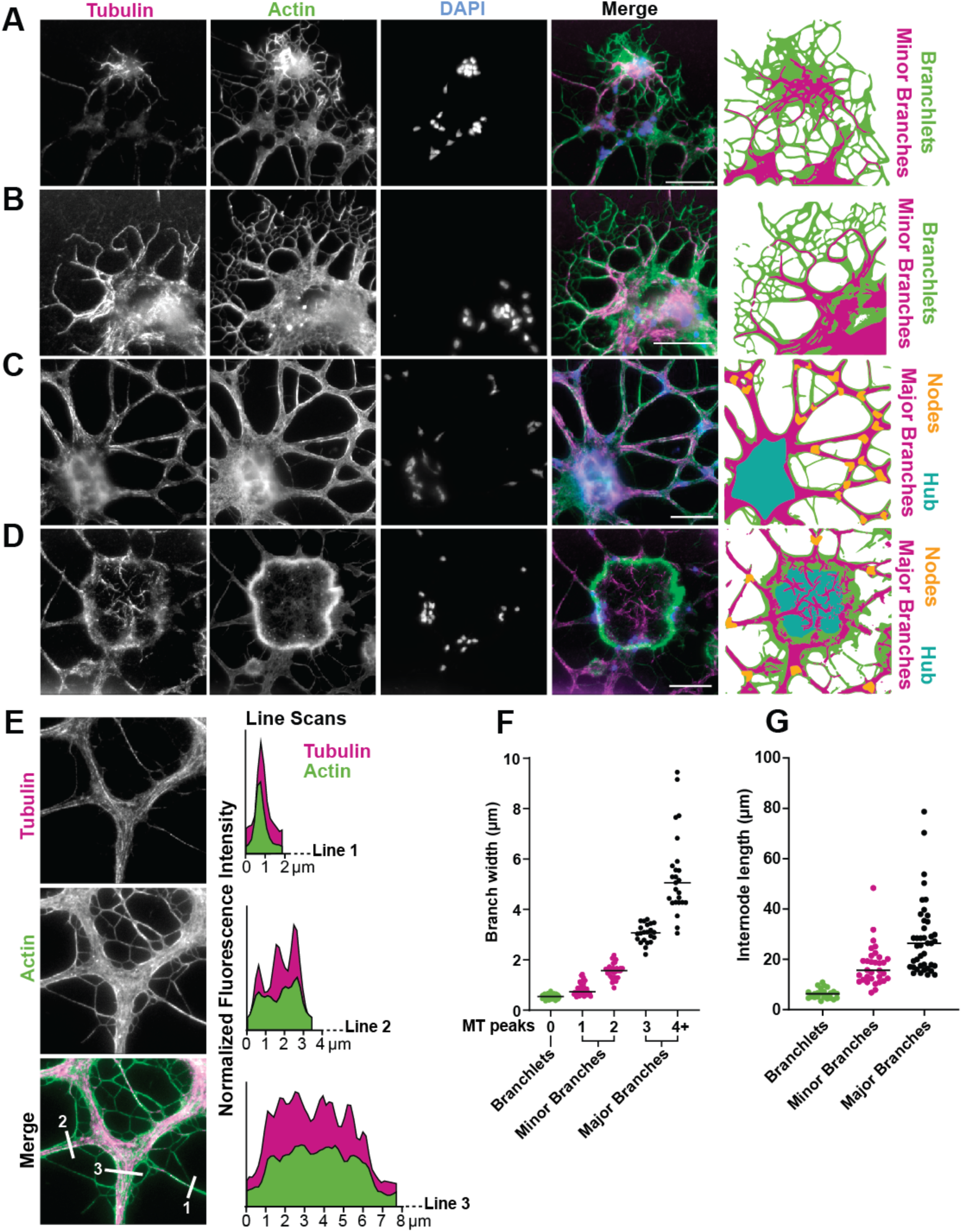
*Filoreta* syncytia have distinct cytoskeletal architectures in pseudopodia and branches. Syncytia were fixed and stained for actin (green), microtubules (MTs) (magenta) and DNA/nuclei (blue) Scale = 20µm for all images. (**A**) A small syncytial fragment (top) fused to a larger syncytium (bottom) showing enriched actin at filopodial (branchlet) fusion sites. The right panel shows a cartoon of features and identification of branchlets and minor branches with newly proliferated MTs between syncytia. (**B**) Peripheral branchlets show partial proliferation of MTs in the filopodial branchlets. The right panel shows a cartoon of features highlighting proliferating MTs into anastomosed branchlets at the transition zone. (**C**) The interior branches of the network include both actin and MTs. Actin is found throughout and is enriched at the submembrane cortex (bright margins). The right panel shows a cartoon of features including completed MT loops with MT-based major branches intersecting at nodes (orange). Branchlets (green) can be seen between major branches. (**D**) Central hubs of the network include MT ends and are bordered by an actin-rich boundary. Right panel shows a cartoon of features including hub (blue) with multiple major branches converging, their MT ends visible in thresholded overlay (magenta). (E) Representative image of branchlets, minor and major branches used in quantifications for (F, G). Top panel = Tubulin, Middle panel= Actin, Bottom panel= merged. White bars designate branch cross-sections measured for counting microtubule (MT) peaks (right graphs), width and length. (Branchlets = absent of MTs) and “Branches” (MT-stabilized). (F) Quantifications of branchlet and branch widths per MT peaks counted in line profiles. (G) Quantifications of branchlet and branch lengths by category.

**Figure 3:**
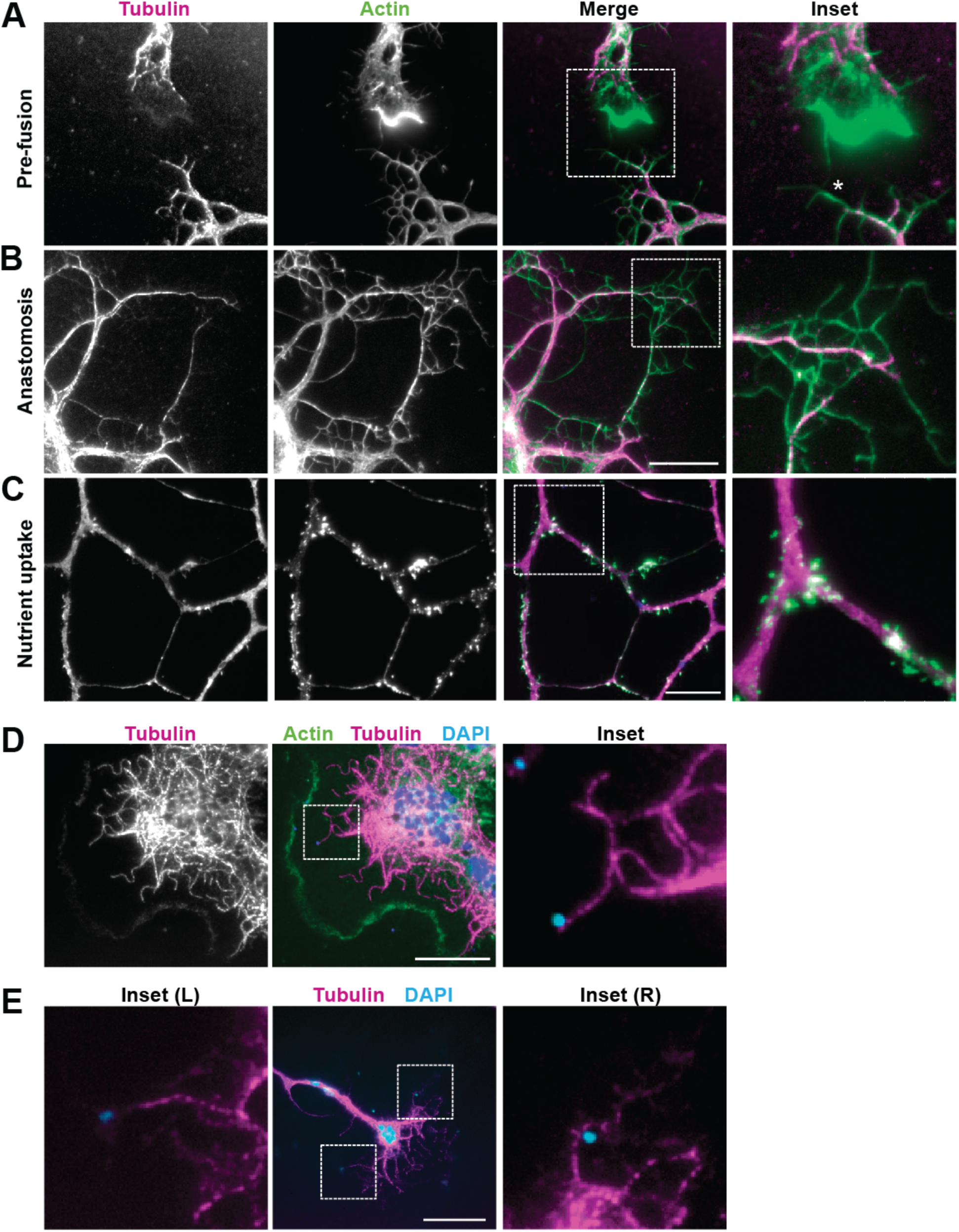
Distinct actin and microtubule structures define syncytial protrusions. Syncytia were fixed and immunostained during specific interactions with their environment (inter-syncytial fusion, self-fusion, increased nutrient availability, and prey capture) with anti-tubulin antibody (magenta) and phalloidin-labeled actin (green). (**A**) Two syncytia exhibiting directional pseudopod extension as they approach fusion. The top syncytium has an actin-rich lamellopod-like protrusion, and the bottom syncytium has numerous filopodia. The site of branchlet (filopodial) fusion is marked with an asterisk. Scale = 20µm. (**B**) Internal branches react to nutrient addition with actin patches within 5 minutes after increased nutrient availability. Scale = 20µm. (**C**) Syncytia branches at the periphery exhibit actin-rich filopodia after being fed enriched medium. Inset shows new reticulopodia at tips of branches. Scale = 50µm. (**D**) A wide lamellipodium with microtubules exhibiting pronounced curvature. The right panel (inset) shows interaction with captured bacterial prey (cyan). (**E**) A branch that has captured several bacteria exhibits straightened microtubules. Scale = 20µm.

To clarify and understand branch development, we next defined characteristics to distinguish “branchlets” from “branches.” Branches leading to hubs appeared 2-3X thicker on average than the peripheral branches closest to branchlets at the periphery. To quantify this impression, we measured the cytoskeletal and morphological aspects of syncytial branchlets and branches, including widths, lengths, and actin and microtubule content in cross sections. We then further distinguished distinct classes based on morphological characteristics. While the widths of branchlets containing only actin varied from 0.3 μm to 0.6 μm, even the narrowest microtubule-filled branches were 24 percent thicker on average (**Figure 2E, 2F**). We defined these as “minor branches.” The branches in the interior of syncytia contained more tubulin signal in distinct quantifiable peaks, which we used to classify minor and major branches. Minor branches contained one or two tubulin peaks, and major branches contained three or more. The branchlets and branch types also varied with respect to length as well. Actin-filled branchlets, like dendritic filopodia (26), were limited to less than 10 μm in length. In contrast, those that included microtubules varied widely, with major branches at least 10 µm and often over 50 μm (**Figure 2G**). This distinction, combined with live observations of transient filopodia that can collapse or retract, suggests that microtubule proliferation into filopodial branchlets stabilizes them for development into branches of the overall network structure. The progressive enrichment of microtubules within thicker and longer-lived branches suggested that branch maturation may depend on selective stabilization through microtubule incorporation. We therefore next examined how distinct actin and microtubule architectures contribute to protrusion dynamics during network growth.

### Distinct actin and microtubule structures define syncytial protrusions

To define the specific cytoskeletal architectures underlying various pseudopods through different stages of development, we fixed and stained syncytia in each stage to visualize the actin and microtubule organization. During neighbor fusion, syncytia in close proximity developed pseudopodial extensions that reach to and fuse with each other, including actin-rich ruffled lamellar pseudopods and branching filopodia (**Figure 3A**). After treating *Filoreta* syncytia with enriched media, established branches developed new lateral branchlets (**Figure 1C, 1I**). Upon staining, we observed dense actin patches along branches, and actin-filled pseudopodial projections that extended laterally from branches. Several of these projections were also enriched with tubulin signal, which is consistent with our prior observation that pseudopodial branchlets initiate new branches in the syncytial interior (**Figure 3B**). To further test the dynamic nature of lateral branching, we treated starved syncytia that were undergoing pruning with enriched medium for 5 minutes prior to fixation. In treated syncytia, peripheral branches were simplified and elongated but had numerous small filopodial branchlets (**Figure 3C**). These observations indicate that actin-dense patches are formed as a response to external stimuli like food.

Microtubule orientation guides and regulates neuronal growth cone turning and migrating cell directionality (27,28). To determine whether microtubules played a similar role in *Filoreta’s* lamellar protrusions, we examined the microtubule structure within lamellar extensions at the growing periphery. We examined syncytia exhibiting lamellar branchlets and saw that they contained microtubules with varying degrees of curvature (**Figure 3D**). To quantify the relationship of microtubule curvature and proximity to actin-rich lamellar regions, we measured each microtubule curvature radius (R_c_) and its distance from the leading edge. Curved microtubules had a R_c_ averaging 1.07 µm ± 0.28 with curve tangents more than 5 µm from the lamellipodial edge. In contrast, straightened microtubules had an average R_c_ over 3.0 µm and were less than 5 µm from the edge. The proximity to the leading edge and loss of curvature suggests these microtubules may have become captured through other protein interactions, and were subsequently straightened under tension, similar to microtubules driving growth cone turning (29). We also observed that straightened microtubules oriented their ends towards phagocytosed bacterial cells (**Figure 3D, 3E**), suggesting a role in prey capture and phagosome transport, as observed in metazoan cell types (30). These observations indicated that exploratory protrusions, branch guidance, and environmental responses are tightly coupled to localized actin–microtubule organization. We next sought to determine how these cytoskeletal interactions shape network topology during syncytial development and remodeling.

### Syncytia undergo dynamic branch remodeling throughout development

The topology of a branched network underscores the direct connection between cytoskeletal structure and its function in efficient transport. During development, neurites extend their branches to grow, make decisions for their patterning based on external cues, and prune to simplify connections throughout maturation (31,32). These developmental features can be morphometrically defined through measurements of branch length and number, enabling better assessment of developmental timepoints and hallmarks for neuron development (33). We therefore sought to define branch patterning throughout network development in a similar manner for *Filoreta.* To quantify the degree of branch arborization in growing syncytia during development, we used Sholl Analysis to measure branch distribution and density at distinct timepoints of *Filoreta*’s life cycle. Sholl Analysis has long been used to quantify the complexity of arborization patterns in neuron morphotypes (34), which defines an arbor’s level of branching complexity as the number of intersecting branch points plotted across radial distances from a central node. Sholl analysis reveals the spatial distribution and density of branches for arborized cells, enabling consistent quantification of morphotype complexity.

To quantify *Filoreta’s* network development during network expansion (**Figure 4A**) and branch extension (**Figure 4B**), we generated Sholl profiles for growing syncytia over 20 minutes and used the resultant graphs to calculate the outward growth rate of the network. Under normal growth conditions, syncytial outgrowth averaged 2.5 μm/minute (± 0.55 μm) (**Figure 4E**), with no significant difference in outward growth rate between radially symmetrical syncytia, and individual branches. To assess if this outward growth coincided with a change in overall branch density, we measured changes in the area under the curve (AUC) as a proxy for surface area covered by the branch intersections. We determined that syncytia and branches expand in surface area coverage by an average of 2.4% per minute at peak growth conditions (**Figure 4F**). This indicates that while the network expanded, it did not do so by eliminating branches and redistributing them; rather, it elaborated on existing branches at the periphery.

**Figure 4:**
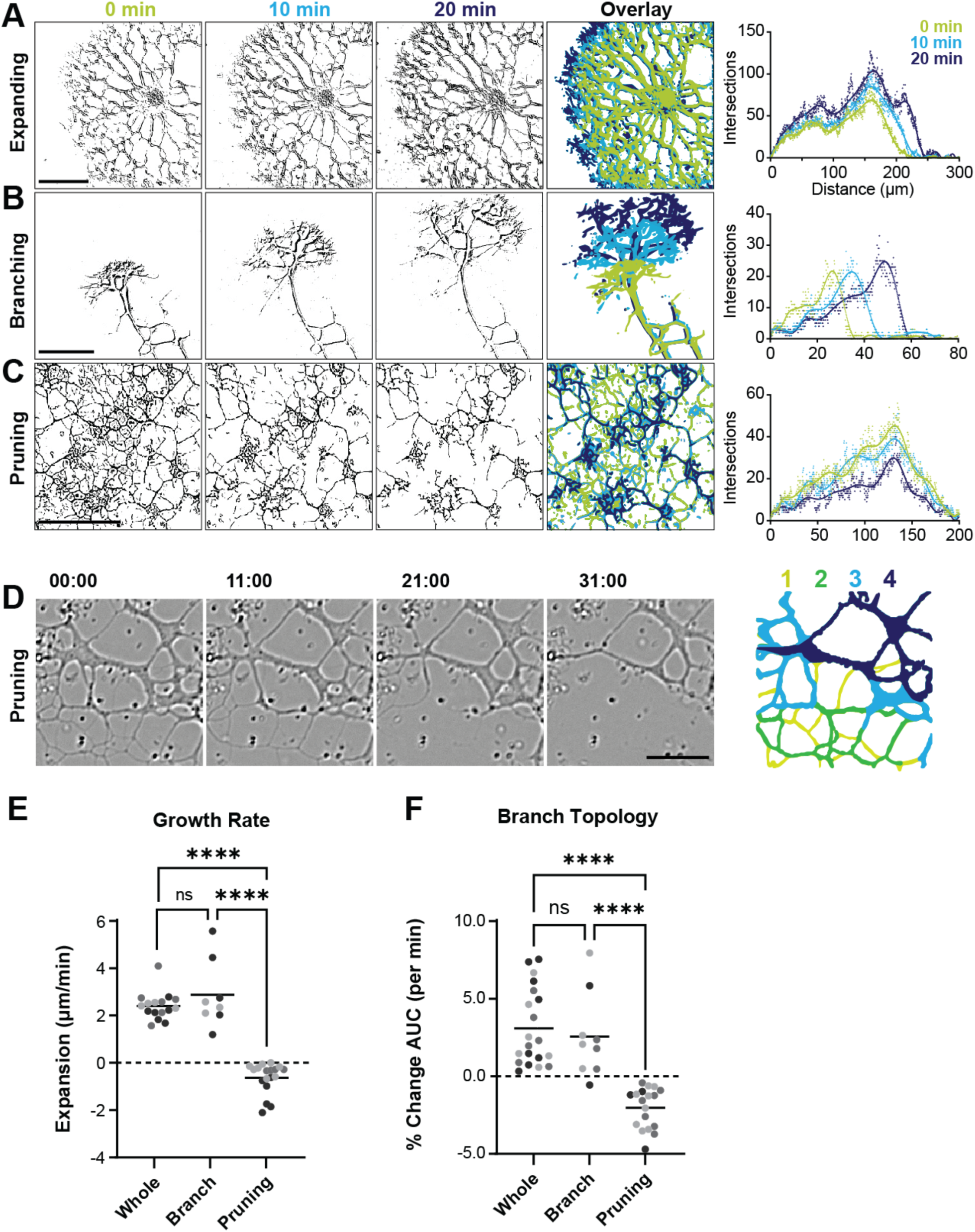
Throughout development, the syncytium undergoes dynamic branch remodeling. Thresholded images showing syncytial branching and their associated Sholl Intersection Profiles (SIPs) at each time point. Time points are color coded chronologically: lime =0 minutes, light blue = 10 minutes, dark blue = 20 minutes. (**A**) The outward growth of a radially symmetrical syncytium over 10-minute intervals. Scale = 100μm. (**B**) Extension of a syncytium with broken symmetry showing elongating branch with arborizing branchlets. Scale = 50μm. (**C**) The entire syncytial network undergoes pruning when exposed to blinking light. Scale = 100μm. (**D**) The order of branch retraction during pruning, with the branch pruning false-colored in the right panel. Branchlets in panel 1 disappear first (chartreuse), followed by minor branches in panel 2 (green), followed by thinning of major branches in panel 3 (light blue) and loss of major branches (final panel, dark blue). scale bar = 20 µm. (**E**) Quantification of peripheral growth rates on individual syncytia from Sholl profiles and (**F**) change in branch topology (percent change in Area Under Curve). n = 8 (branch) n = 16 (whole) n = 18 (pruning). ****p<0.0001, by ANOVA. Data sets colored per replicate.

We next wanted to assess the changes in topology during pruning, which occurs at varying rates depending on the severity of the environmental factors affecting a syncytium (**Figure 4C**). For these experiments, pruning was induced through removal of nutrient medium and exposure to blinking light during imaging (see Methods). Stressed syncytia retracted quickly, with approximately 1.5% loss of surface area coverage per minute (**Figure 4F**). Syncytia simplified arbors by pruning pseudopodia and branchlets in a manner that first narrowed minor branches and then major branches, until branches retracted to hubs (**Figure 4D**). Together, these measurements provide a basis to quantify the dynamics of syncytial development and enable assessment of network complexity. The ability to quantify branch elaboration and pruning allowed us to establish a framework for testing how actin and microtubule systems contribute to network growth, maintenance, and remodeling.

### The dynamics and growth in syncytial arbors depend on actin and microtubule polymers

Based on our live imaging observations of branchlet transience, and the microtubule content in interior branches, we hypothesized that microtubules proliferate into branchlets and stabilize them. Having defined the stages of network development and quantified the rates of outgrowth in normal conditions, we next sought to define the respective roles of the microtubule and actin cytoskeleton in generating the arborized morphology. To test these cytoskeletal components independently, we treated growing syncytia with the microtubule polymerization-interfering drug nocodazole and the F-actin disrupting drug Latrunculin A. As an internal control for variability in culture replicates, we quantified syncytial growth and distribution before (**Figure 5A, 5C**) and after each treatment (**Figure 5B, D**). Sholl profiles generated for each treatment period showed unique changes in branching for the different cytoskeletal targets. Following nocodazole treatment, the expansion rate dropped to 0.4 μm/minute, while branch density decreased as seen during pruning (**Figure 5E, 5F)**. Some branches retracted to proximal nodes, suggesting microtubule depolymerization toward the minus ends, loss of branch stabilization, or both (**Figure 5G**). This pattern also implied the existence of arrays of antiparallel-bundled microtubules. Longer term drug treatment (40 minutes) caused truncated branches and loss of long-range syncytial contacts, yet growth was recoverable upon nocodazole washout and media replacement (**Video S2**). This indicates microtubule proliferation into branchlets during outgrowth plays a role in branch stability, and that microtubules likely provide structural integrity for interior branches.

**Figure 5:**
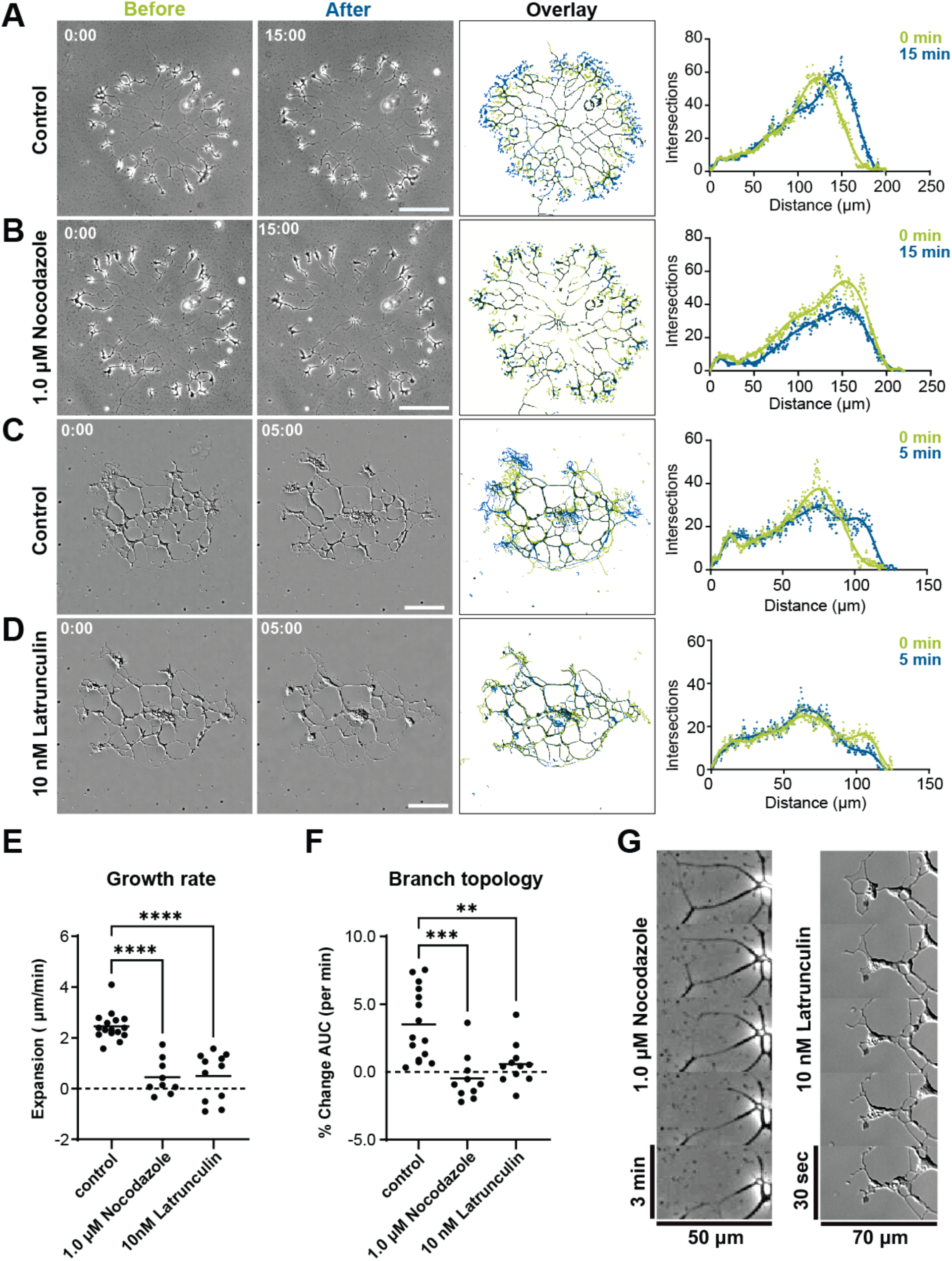
Syncytial arbors are dependent on actin and microtubule polymers. (**A-B**) A growing syncytium was imaged for 15 minutes before (**A**) and after (**B**) treatment with 1.0 µM nocodazole. The growth over 15 minutes is visualized by a false-colored thresholded overlay (3rd panel) showing each timepoint superimposed (lime = 0 minutes, blue = 15 minutes, black = combined; locations branches remained unchanged between both images). Scale bar = 100µm. The far-right panels show the SIP for each timepoint. (**C-D**) A growing syncytium was imaged for 5 minutes before (**C**) and after (**D**) treatment with 10nM Latrunculin A. Scale bar = 50µm. The images were thresholded as above and SIPs are shown for each timepoint. (**E**) The growth rate of syncytia is significantly affected by cytoskeletal drugs. These experiments were repeated on different syncytia to quantify effects of each drug on the syncytial arbors. The growth rate of each syncytium was calculated by the difference in SIP distances as µm/minute. (**F**) Branch topology is significantly affected by drug treatments. The area under the curve (AUC) was calculated between time points and presented as a change in percentage of AUC at the starting point. The growth rate and change in AUC were statistically compared using ANOVA (**p=0.0017; ****p<0.0001). Control (n=19), Nocodazole (n=11), Latrunculin (n=11). (**G**) Montages of syncytial branches during each drug treatment, showing internal branch loss during Nocodazole treatment (left panel) and peripheral branchlet retraction during Latrunculin A treatment (right panel).

We then queried the actin-dependence of branch initiation and network complexity by examining treatments with Latrunculin A (**Figure 5C, 5D**). In contrast to the internal controls’ normal outgrowth in the same time span, we saw that Latrunculin-treated syncytia ceased peripheral branching, and many filopodia and lamellipodia seized and retracted (**Figure 5G**). Branchlet extension decreased overall, while interior branch topology was largely retained (**Figure 5F**). This indicates actin has an important role for branchlet development, but its polymerization is not integral to the maintenance of branches once they are established. Selective stabilization of microtubule-enriched branches following actin disruption indicated that branch maintenance becomes increasingly microtubule-dependent during network maturation. We therefore next quantified how cytoskeletal perturbations alter the relative organization of actin and microtubule networks throughout the syncytium.

### Nocodazole significantly alters the microtubule distribution throughout the network

Having determined that both nocodazole and Latrunculin A disrupt network development, we wanted to visualize the actin and microtubule distribution and quantify the underlying changes to the polymers during these drug treatments. To directly compare the relative distributions of microtubules in internal and peripheral regions of the network before and after drug treatments, we treated syncytia with nocodazole, then fixed and stained the resultant networks. We then performed Sholl analyses on the actin and microtubule fluorescent signals separately to quantify the microtubule network as a percentage of the actin network. In Sholl profiles from untreated syncytia (**Figure 6A, 6C**), the interior branches had 88.15% (± 4.9%) microtubule staining, while branches in the periphery had only 65.55% (± 5.4%). We then compared these measurements to the relative distribution of microtubules and actin in nocodazole-treated syncytia (**Figure 6B, 6D**). At the interior, microtubule coverage dropped to 65.36% (25% decrease), where the periphery dropped to 38.76% (40% decrease) (**Figure 6F**). This suggests that the microtubules at the periphery are more sensitive to nocodazole, potentially by lack of microtubule -bundling, binding and stabilizing proteins, or more dynamic instability during proliferation.

**Figure 6:**
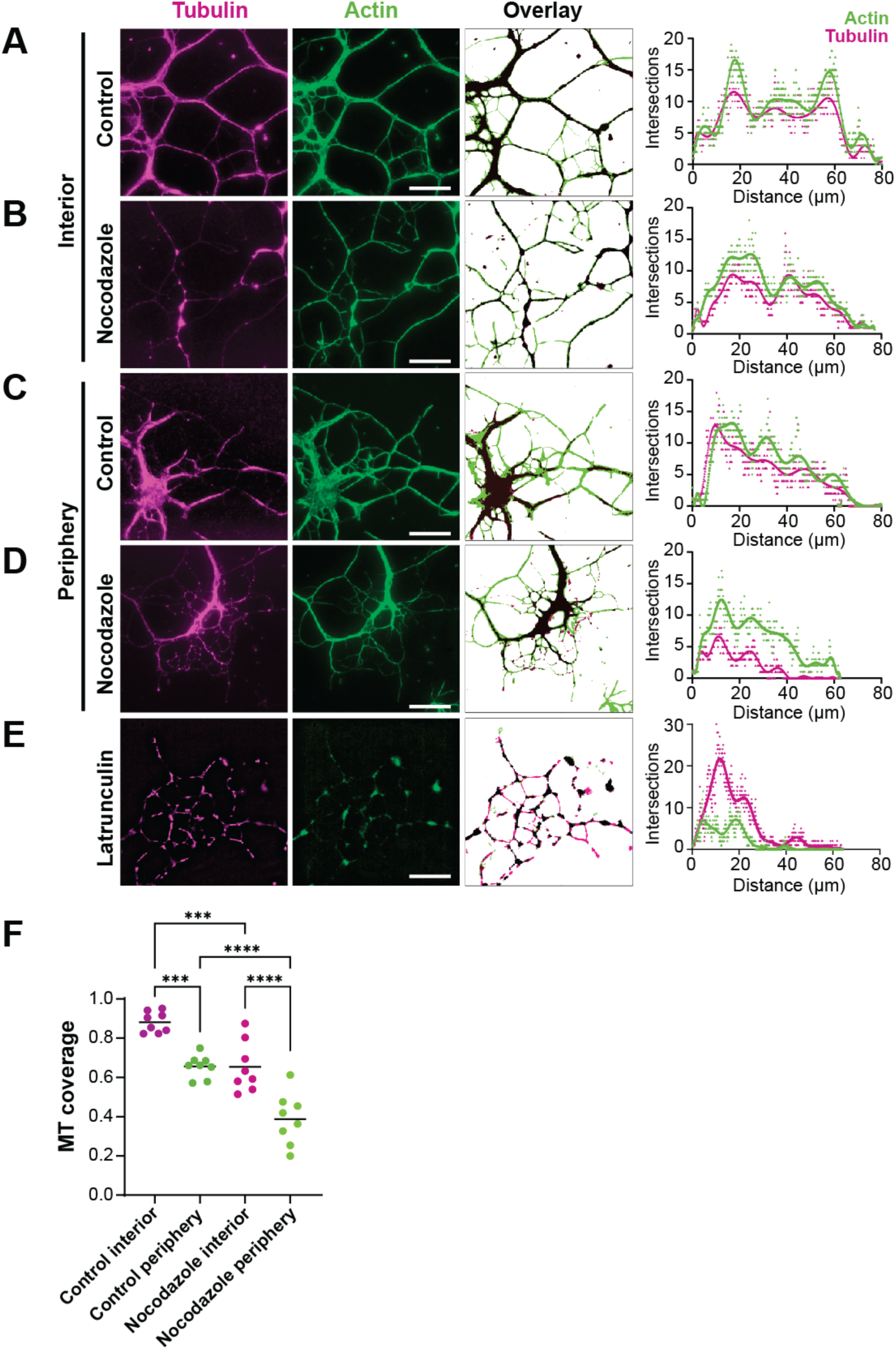
Microtubule distribution and branch stability is variable between the network interior and periphery. Syncytia were fixed and stained under normal conditions or after treatment with 1.0 µM Nocodazole. (**A**) The branch topology of the syncytial network interior, stained for microtubules (magenta) and actin (green). A false-colored overlay (third panel) shows the proportion of microtubule -filled branches (microtubule = magenta, actin = green, both = black) and branchlets (actin only, green). The corresponding SIP (far right) for each cytoskeletal component’s quantified distribution. (**B**) The microtubule and actin distribution of a nocodazole-treated syncytium interior. The false-colored overlay shows tubulin staining relative to actin-stained branches, with SIP shown as before. (**C-D**) The same staining and thresholding technique was repeated for the periphery of untreated syncytia (**C**) or nocodazole-treated syncytia (**D**). (**E**) A syncytium treated with latrunculin has altered distribution of microtubules and actin. The false colored overlay and corresponding SIP shows relative distribution for actin and microtubules. Scale = 20 µm for all images. (**F**) Relative cytoskeletal SIPs for nocodazole treatments, quantified as a percent of microtubule coverage within the actin-stained network. The latrunculin treatment shown in (**E**) was excluded, as the actin was not measurable to use for the total area covered by the arbor. The interior and periphery of untreated syncytia have significantly different microtubule distributions (*** p=0.0006). The microtubule distribution is significantly different between both interior (***p=0.0005) and periphery of nocodazole-treated syncytia, compared to untreated (****p<0.0001), by ANOVA. n=8 for all categories.

To test whether actin drugs affect microtubule distribution, we treated syncytia with Latrunculin and made the same comparison (**Figure 6E**). We saw that although they appeared somewhat punctate, the topology of microtubules extended beyond the actin topology, indicating the microtubule distribution remained unchanged even in the absence of polymerized actin with the branches and branchlets. Taken together, these data support our hypothesis that microtubule proliferation of branchlets at the periphery leads to their stabilization, and that branches that include more microtubules are more stable than branchlets that are partially filled.

### Organelles are rapidly and bidirectionally transported throughout the syncytial network

Rapid organelle transport is a feature seen in other rhizarian amoebae, like *Reticulomyxa filosa* (35,36). Our preliminary live imaging indicated the *Filoreta* syncytium also undergoes rapid organelle transport. To further investigate this phenomenon, we visualized organelle movements during live imaging by staining established syncytia with Hoechst 33342 (nuclei), Syto9 (mitochondria), and Lysotracker (lysosomes). Nuclei, mitochondria, and lysosomes were transported rapidly and bidirectionally throughout the branches of the syncytial network, at rates averaging 5.3 µm/sec, 8.3 µm/sec, and 7.3 µm/sec, respectively (**Figure 7A, 7B**). Organelles traveling within the same branch were not constrained to the same rate or direction (**Figure 7C**). Additionally, the transport rate often slowed or stalled as organelles enter nodes, where two or more branches intersect (**Figure 7D**). We hypothesized that branch width is related to rate of organelle transport. To quantify this relationship, we generated kymographs of organelle movement through a variety of branch widths (1.5 µm to 6.5 µm). We determined that the rate of organelle transport loosely correlated with the width of the branchlet (R^2^ = 0.5498) (**Figure 7E**). For uncomplicated branches (i.e., fewer than one node every 10 µm in length), the rate of transport generally increased with branch width. This matched our prior observations showing that within the same branches, transport was not a constant rate for all organellar cargoes. However, the approximate number of microtubules in minor and major branches does directly correlate with branch width, (R^2^ = 0.9029) (**Figure 7F, 7G**). Combined, these data suggest that longitudinal microtubules are used as a means of organelle transport. Additionally, branch thickness may be an important factor regulating organelle transport in different regions of syncytia and during different stages of development. The strong relationship between branch width, microtubule abundance, and organelle transport suggested that syncytial network architecture may scale with the degree of intracellular transport.

**Figure 7:**
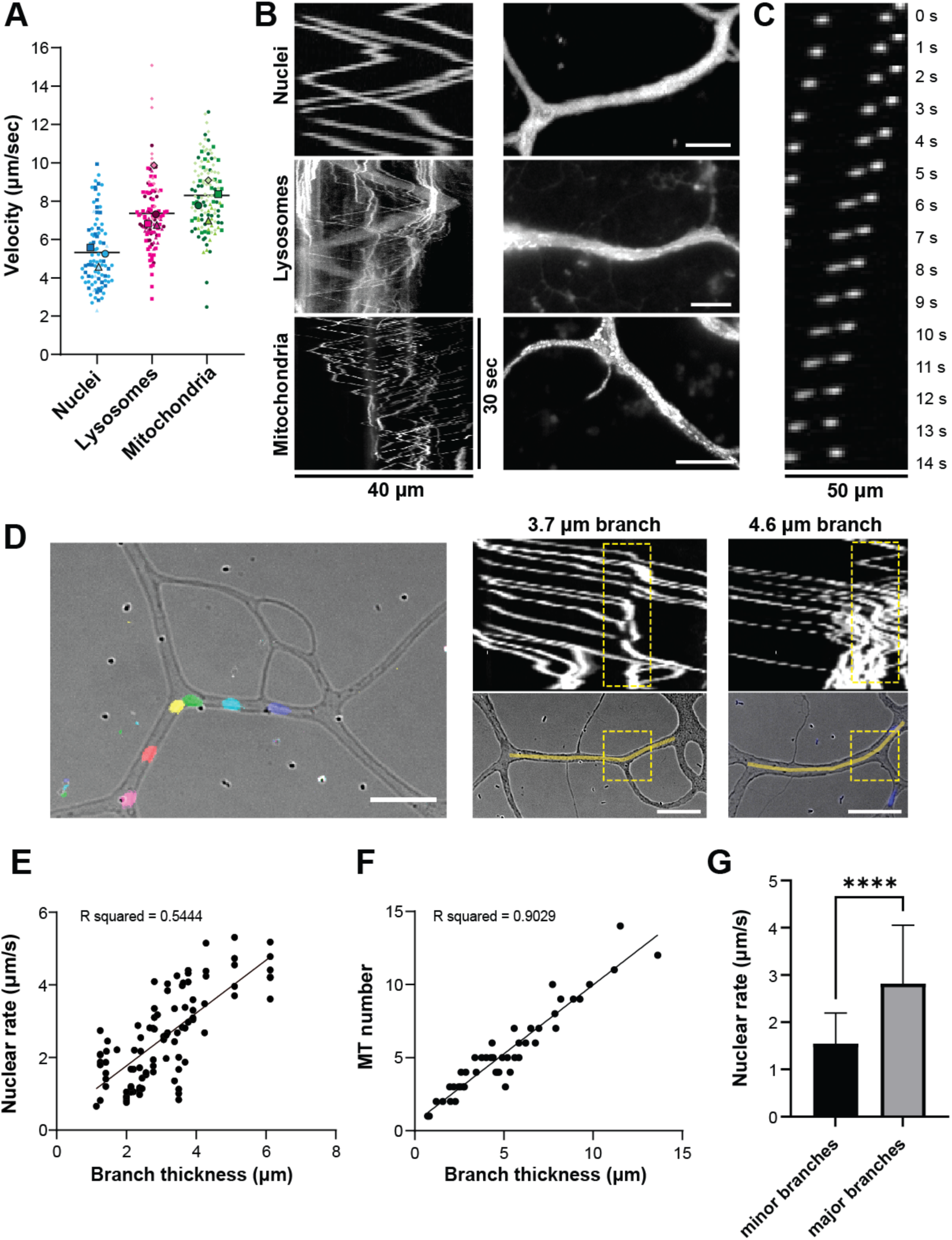
Organelles are rapidly and bidirectionally transported throughout the syncytial network. (**A**) Syncytia were stained separately with organelle markers: Hoechst 33342 (nuclei), Syto9 (mitochondria), and Lysotracker (lysosomes), and their respective rates were quantified. Datasets colored per technical replicate; nuclei (3; n=82), lysosomes (4; n=89) mitochondria (4; n=55). (**B**) Representative kymographs of organelle movements over 40μm and 30 seconds. Right panels show branches used for kymographs on the left, created as stack overlays. Scale=10 μm. (**C**) Organelles within the same branch move independently. A montage shows nuclei are transported in a branch at different rates. The nucleus on the right changes direction at T=13s. (**D**) Nuclei traveling in branches become stalled or slow at nodes. The left panel is false-colored with 5 second intervals, scale = 20μm. The right panels show kymographs of nuclear movements corresponding to the bottom panel branch images. The locations of branch nodes in the kymographs are outlined in yellow and correspond to inset squares. (**E**) The nuclear rate plotted as a function of branch thickness, R-squared = 0.544. (**F**) Microtubule number, approximated by peaks in line scans, plotted with branch thickness. R-squared = 0.903. (**G**) The transport rate quantified in (**E**) organized into minor and major branch categories based on branch thickness (minor branches <2.5 μm, major branches >2.5μm). ****p<0.0001 by unpaired T-test. (n=19 minor branches, n=76 major branches).

### Nuclei and other organelles are transported throughout the syncytial network using microtubule-dependent motors

Organelles traveling within the same branch simultaneously move at different rates and directions, inconsistent with cytoplasmic streaming as the primary force for long-ranged transport. To test if organelles are instead transported by microtubules, we treated Hoechst 33342-stained syncytia with nocodazole to track changes in nuclear transport during live imaging (**Figure 8A, B**). While nuclei were still capable of bidirectional movements, their translocation distance decreased by 33%, and maximal rate by 27% for 20 μm nocodazole treatments. Nuclear transport was most diminished at major branch nodes, where nuclei stalled completely (**Figure 8E**). This indicated that nuclei and other organelles travel along microtubules, which likely originate at nodes.

**Figure 8:**
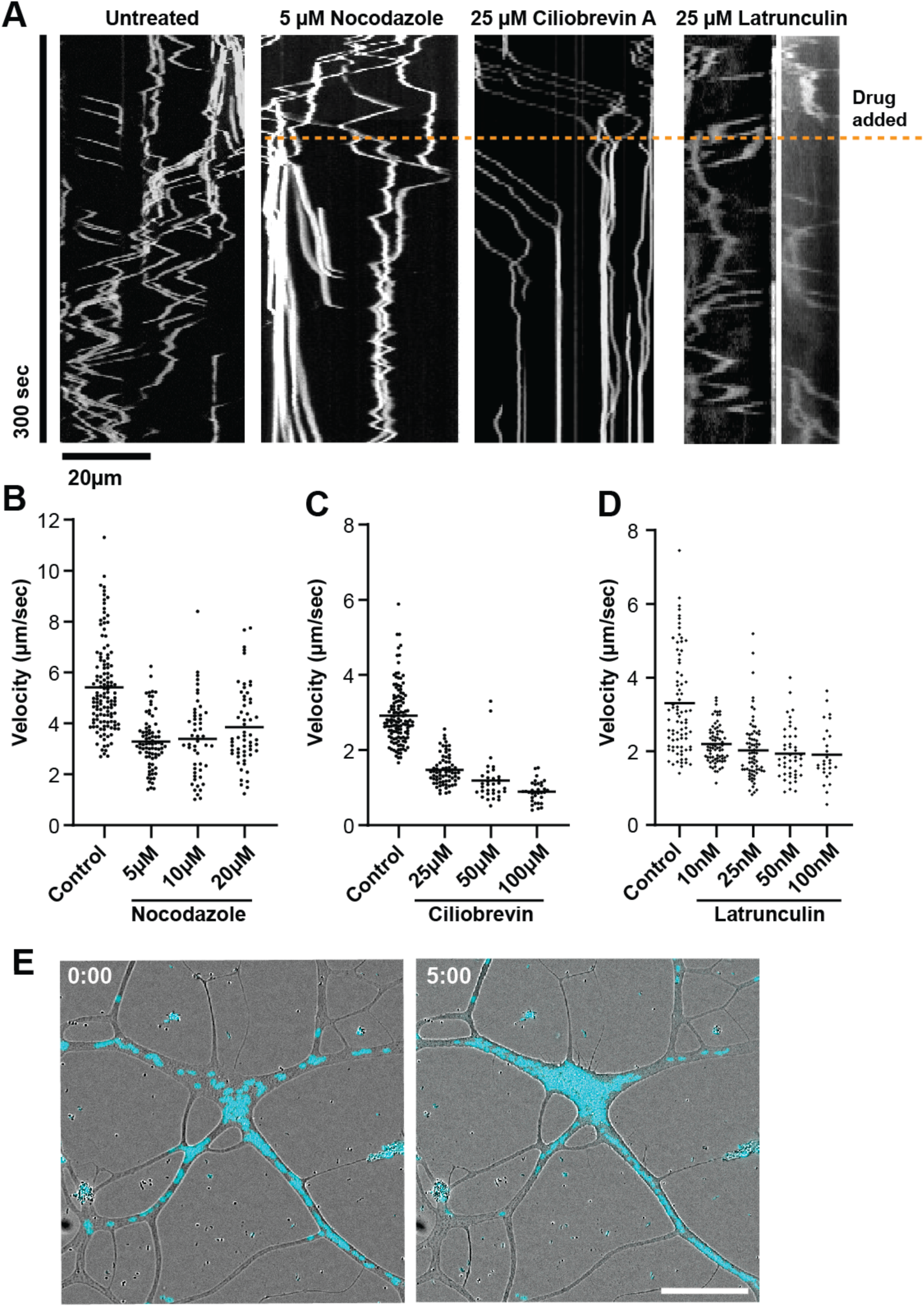
Nuclei are transported throughout the syncytial network using microtubule-dependent motors. Syncytia were stained with Hoechst 33342 and imaged during various drug treatments (Nocodazole, Ciliobrevin D, Latrunculin). The respective cytoskeletal drugs were added at the 1-minute timepoint (dotted line). (**A**) Representative kymographs showing nuclear movements over a 5-minute time course and were treated with the specified cytoskeletal drug concentration at the indicated timepoint, marked with a dotted yellow line. (**B-D**) Nuclear rates were quantified in μm/second for processive movements of 10μm or longer. (**B**) Rates measured in treatments with nocodazole concentrations of 5 μM, 10 μM, and 20 μM. (**C**) Ciliobrevin D treatments measured at concentrations of 25 μM, 50 μM, and 100 μM. (**D**) Latrunculin treatments measured at concentrations of 10nM, 25 nM, 50 nM, and 100 nM. All treatments were statistically compared using one-way ANOVA against the buffer control. (**E**) Effect of 20 µM nocodazole on nuclear trafficking at t=0 (left panel) and t=5 min (right panel) (nuclei = cyan). Scale = 50 μm.

Following the notion that depolymerization of microtubules results in shortened “tracks” for nuclear translocation, eventually trapping nuclei in nodes, we surmised that microtubule nucleation likely occurs at nodes, or that their minus ends are stationed there. A distant rhizarian relative, *Reticulomyxa filosa*, has an extensive repertoire of minus-end directed microtubule motor proteins called dynein (37), including a unique dynein capable of bidirectional organelle transport (36). To determine whether *Filoreta* also transports its organelles using dynein motors, we treated live syncytia with the dynein inhibitor Ciliobrevin D and imaged nuclear movement (**Figure 8A**). After treatment with Ciliobrevin D, there was an immediate and significant reduction in both the rate and distance of nuclear movement (**Figure 8C**). Over 50% of the nuclei moved processively (more than 10 µm) before treatment, and only 6.5% moved processively after addition of 100 µM Ciliobrevin. Of the nuclei that still moved, their rates were slowed by 70%. However, the directionality of the remaining mobile nuclei was not significantly affected, and many nuclei still moved bidirectionally through branches at the lowest Ciliobrevin concentrations. While this could be consistent with a Ciliobrevin-resistant bidirectional motor, we could not exclude the possibility that actin-based forces also contribute to organelle transport. To investigate this further, we treated syncytia with a range of concentrations of the F-actin inhibitor Latrunculin (10, 25, 50, 100 nM). Latrunculin treatments had a slightly decreased rate of transport (< 25% difference) (**Figure 8D**), but the more apparent effect of the drug was its impact on the morphology of branches as concentrations increased (**Figure 5D**). Together, these results suggest that dynein is the major contributor to organelle translocation along microtubules, and actin plays a supporting role. Branch nodes may function as structural intersections and organizational centers for microtubule arrays and new nodal pathways for intracellular transport.

### Microtubule nucleating complexes localize to branch nodes independent of nuclear location

The observed patterns of microtubule arrays, combined with nuclear trafficking and stalling in nodes after nocodazole treatments, was consistent with branch nodes as the primary site of microtubule nucleation. A common mechanism for microtubule nucleation uses gamma-tubulin ring complexes (ɣ-TURC), which initiate microtubule polymerization at specific sites within cells. These sites include nuclear-associated microtubule organizing centers (MTOCs) but can also initiate polymerization laterally along existing microtubules in a process called branched microtubule nucleation (38). We hypothesized that ɣ-TURC localizes to nodes of bifurcating and anastomosing branches in developing syncytia, which enables further elaboration of the network and extension of microtubule tracks for organelle distribution. To visualize the sites where the microtubule network is nucleated and organized, we stained fixed syncytia with an AlexaFluor conjugated gamma-tubulin antibody (**Figure 9A**). Gamma tubulin staining was diffuse throughout the microtubule network but had robustly stained puncta in branch nodes (**Figure 9B**). To confirm the gamma tubulin localization, we additionally raised antibodies specifically to the *Filoreta* GCP3 sequence, a component of ɣ-TURC (39). The GCP3 staining colocalized with gamma-tubulin (**Figure 9E**), and was enriched in nodes (**Figure 9C, D**). In the syncytial hubs which contained numerous nuclei, we observed that gamma tubulin and GCP3 puncta colocalized but did not coincide with nuclear location (**Figure 9E**). In the larger branches emanating from hubs, both GCP3 and gamma-tubulin colocalized with each other in puncta at the node junctions. This indicates that microtubule organization is not centralized around nuclear location like canonical MTOCs and suggests that the extensive microtubule array is self-elaborating through branched nucleation machinery.

**Figure 9:**
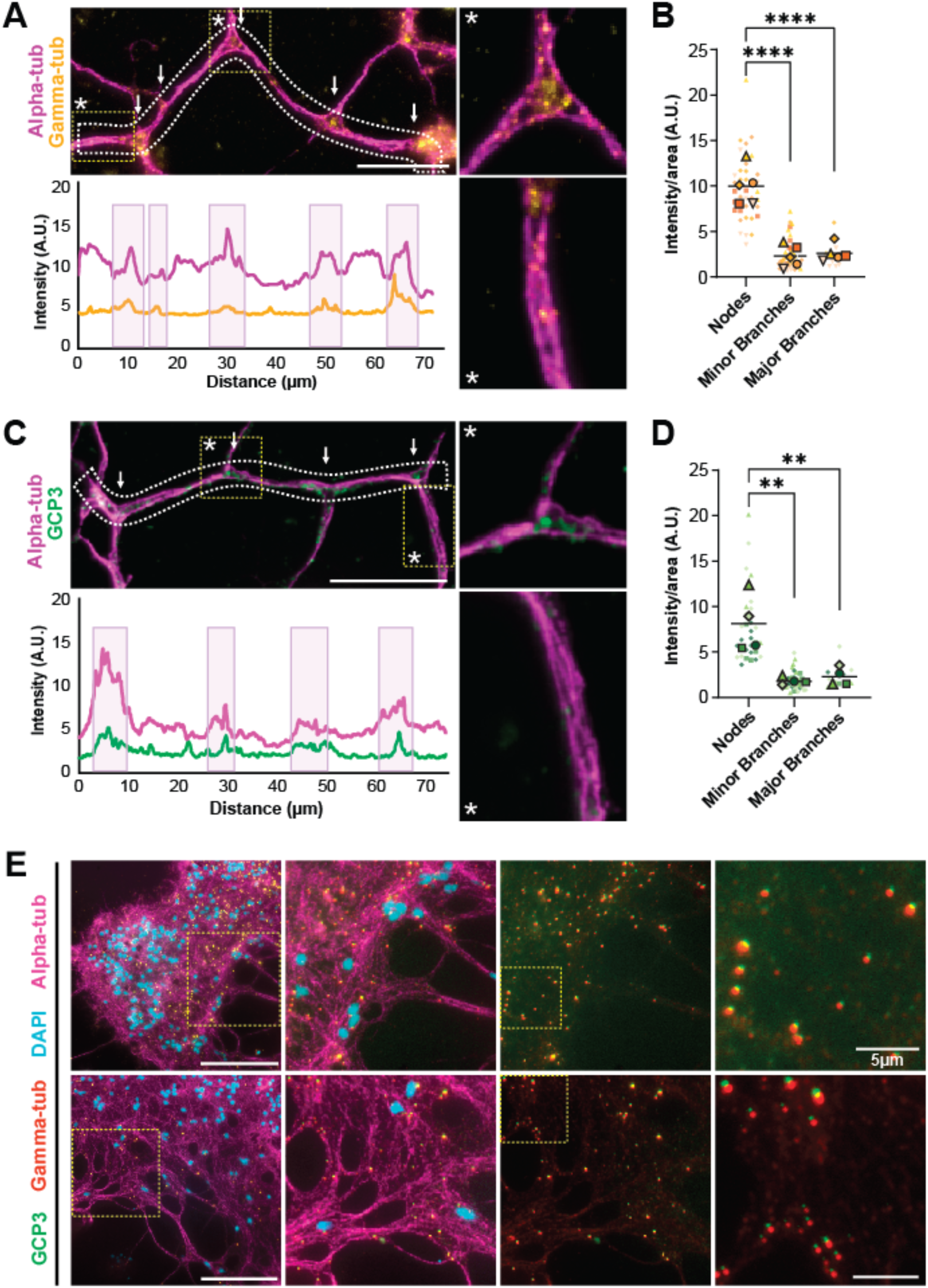
Microtubule nucleating complexes localize to branch nodes independent of nuclear location. (**A**) Gamma-tubulin (yellow) foci are enriched at branch nodes (shown by alpha-tubulin, magenta) in fixed and immunostained syncytia. Intensity profiles are shown for areas within 50-pixel wide line scans (dotted lines). Regions surrounding nodes (white arrows in image, shaded region in graph) have increased tubulin signal due to additional microtubules in that region. The gamma-tubulin signal has distinct peaks within nodes. Insets (right) marked by asterisks show a closer view of a node (top) and branch proximal to a node (bottom). (**B**) Quantification of the signal intensity per pixel area for gamma-tubulin in nodes vs minor and major branches, measured across 5 replicates. Statistical significance by ANOVA (**C**) The gamma-tubulin ring complex component GCP3 (green) shows similar patterning in fixed and immunostained syncytia. Intensity profiles, inset images shown as described above for gamma-tubulin. (**D**) Quantification of the signal intensity per pixel area for GCP3 in nodes vs minor and major branches, measured over 4 replicates. Statistical significance by ANOVA. (**E**) Two representative sets of images of syncytia stained with conjugated gamma-tubulin antibody (red), custom GCP3 antibody (green), alpha-tubulin (magenta) and DAPI (blue). Far left: A broad syncytial hub with nuclei. Scale bar = 50 μm. The first inset marked by a dotted line is shown in the 2nd column. Images in the 3rd column to the right show the same image as column 2, with only the GCP3 and gamma-tubulin signal. The last column (far right) shows an inset of the 3rd column. Scale bar = 5 μm.

## DISCUSSION

### Conserved cytoskeletal mechanisms underlying arborized network organization across diverse eukaryotic lineages

The development of the *Filoreta* syncytium reveals a hierarchical relationship between actin-driven exploration and microtubule-dependent stabilization during arbor formation. Actin-rich branchlets continuously probe new cellular territory, but only a subset become incorporated into the mature network through microtubule recruitment and stabilization (Figures 2, 5, and 6). Branches that acquire longitudinal microtubule bundles persist and support intracellular transport, whereas branches lacking robust microtubule incorporation remain transient. The transition from dynamic branchlets to stable transport-capable branches therefore coincides with progressive microtubule incorporation, linking branch maturation directly to cytoskeletal specialization.

Several aspects of this branching mechanism parallel observations from neuronal arborization. In both systems, actin-rich protrusions precede branch stabilization, while microtubules selectively reinforce persistent branches (Figures 2, 3, 5, and 6). The enrichment of γ-TURC components at branch nodes in *Filoreta* syncytia further indicates distributed microtubule nucleation rather than centralized centrosome-like organization (Figure 9)(40), while the dependence of mature branches on microtubule integrity links branch persistence to microtubule-dependent stabilization despite substantial differences in overall cellular architecture (41,42).

Comparable relationships between exploratory actin structures, microtubule-dependent branch stabilization, distributed nucleation, and intracellular transport have also been described in fungal hyphae, epithelial branching morphogenesis, and amoeboid lineages that generate extensive pseudopodial networks (9,43,44). Classical studies of the reticulose amoeba *Reticulomyxa filosa* similarly demonstrated rapid bidirectional transport along longitudinal microtubule arrays throughout an extensive arborized network (22,23,36), although how such networks were generated and maintained remained unresolved. In *Filoreta*, the combination of cytoskeletal perturbations, morphometric analyses, intracellular transport measurements, and distributed microtubule nucleation markers (Figures 2–9) links branch stabilization, transport, and network architecture within a single mechanistic framework. Together, these observations place the classic transport studies of *Reticulomyxa* into a broader mechanistic framework in which branch formation, transport organization, and network architecture emerge as integrated features of large arborized cells. Viewed through an evolutionary cell biology lens, these parallels are notable not because *Filoreta* resembles neurons, but because arborized systems repeatedly encounter similar scaling constraints.

### Reticulated networks represent an alternative architectural solution to cellular scaling

While neuronal arbors are typically organized as polarized, tree-like structures, *Filoreta* develops a highly interconnected reticulated network through repeated branch fusion and anastomosis. This topology produces a fundamentally different form of arborization in which branches form an integrated syncytial network rather than discrete terminal processes (Figure 1). Similar reticulated architectures have evolved independently in diverse biological systems, including fungal mycelia, slime molds, and vascular networks, where interconnected pathways facilitate transport and communication across large spatial scales (45–47). The presence of an analogous architecture in *Filoreta* suggests that branching can be organized in multiple ways while supporting similar functional demands.

The reticulated organization of *Filoreta* likely provides several advantages for a large syncytial amoeba. Anastomosis creates multiple routes between distant regions of the network, generating redundancy that may increase resilience to local branch loss or disruption (46,48). This topology is accompanied by a decentralized mode of cytoskeletal organization in which nuclei, organelles, and microtubule nucleation sites are distributed throughout the syncytium. The enrichment of γ-tubulin and GCP3 at branch nodes (Figure 9) indicates that microtubule nucleation occurs at multiple locations rather than from a centralized organizing center. Together, these observations suggest that large, arborized cells can maintain connectivity, transport, and network remodeling through distributed cytoskeletal mechanisms embedded throughout the syncytium. Tree-like neuronal arbors and reticulated amoeboid networks therefore demonstrate that similar cytoskeletal mechanisms can generate fundamentally different architectural outcomes. Rather than converging on a single arborized morphology, evolution appears capable of producing distinct topologies that balance transport, connectivity, and network resilience in different ways.

### Cytoskeletal arborization as a recurring evolutionary solution to cellular scaling constraints

The cytoskeletal architecture of *Filoreta* illustrates how branching, active transport, and distributed cytoskeletal organization operate together to support large cellular networks. By linking branch stabilization, transport, and distributed nucleation within a single system, *Filoreta* provides a mechanistic framework for understanding how common scaling constraints can repeatedly shape the evolution of arborized cellular architectures across eukaryotes. Dynamic branchlets increase the area available for environmental sensing, prey capture, and interaction with surrounding substrates (Figures 1–4). However, elaboration of increasingly complex arbors also increases the distances over which nuclei, vesicles, and other organelles must be distributed. Branching therefore couples increased environmental access to increasing demands on intracellular transport and structural coordination. These challenges are characteristic of large cells more generally, where increasing size imposes constraints on transport, communication, and spatial organization (49,50).

The cytoskeletal architecture of *Filoreta* appears adapted to balance these competing requirements. Actin-rich protrusions drive local exploration and branch initiation, whereas longitudinal microtubule arrays stabilize mature branches and support long-range transport (Figures 2, 5–8). Branch elaboration and intracellular logistics therefore emerge as tightly coupled processes rather than independent cellular functions. Selective stabilization of microtubule-containing branches following actin disruption further indicates that branch maturation involves a transition from exploratory growth to transport-capable structural networks.

Several observations indicate that network architecture itself is shaped by transport demands. Organelles move rapidly and bidirectionally throughout the syncytium (Figures 7 and 8), transport rates scale with branch width and microtubule abundance (Figure 7), and nuclei stall at branch intersections following microtubule disruption (Figure 8). Branch nodes therefore function not only as structural intersections but also as transport hubs that coordinate movement throughout the network. The enrichment of γ-tubulin and GCP3 at these sites (Figure 9) further suggests that local microtubule nucleation contributes to maintaining and remodeling the network. Thus, branch formation, transport organization, and network maintenance are not separable processes but emerge as integrated features of large, arborized cells. Distributed organization of this type has been proposed as a mechanism by which large multinucleate cells maintain local control and coordination across extensive cellular networks (49).

The recurrence of arborized architectures across eukaryotes indicates that branching is a common adaptation to the challenges imposed by increasing cellular scale. Neurons, fungal hyphae, slime molds, vascular systems, and reticulose amoebae all employ branching architectures despite profound differences in evolutionary history and cellular organization. Although these systems differ in topology and molecular implementation, each must coordinate transport, environmental interaction, and structural integrity across large spatial domains. Similar scaling constraints therefore repeatedly favor branching architectures and associated transport systems as effective solutions to the challenges imposed by increasing cellular size (50,51). The widespread occurrence of arborized architectures across distantly related eukaryotes indicates that increasing cellular scale repeatedly exposes similar cellular design problems, thereby biasing evolution toward a limited set of successful cytoskeletal solutions. Increasing cellular scale therefore acts not only as a source of selective pressure, but also as a constraint that channels cytoskeletal evolution toward a limited set of successful architectural solutions. The cytoskeletal architecture of *Filoreta* illustrates how branching, active transport, and distributed cytoskeletal organization operate together to support large cellular networks, providing a mechanistic framework for understanding the repeated evolution of arborized cellular architectures across diverse eukaryotic lineages.

### Evolutionary origins of arborized cellular architectures

The hierarchical relationship between actin-rich exploratory branchlets, microtubule-dependent stabilization, intracellular transport, and distributed nucleation observed in *Filoreta* (Figures 2, 5–9) parallels mechanisms described in neuronal and other arborized systems (35,40,52–59). These recurring interactions indicate that common cytoskeletal principles accompany the development of complex arborized architectures across diverse eukaryotic lineages.

Several evolutionary processes could contribute to the repeated emergence of arborized architectures, including deep conservation, repeated co-option of ancestral machinery, and convergence. Deep conservation would imply that fundamental interactions between actin, microtubules, and intracellular transport systems were already present in ancestral eukaryotes and have been retained across lineages (60,61). Alternatively, arborized architectures may reflect repeated co-option of conserved cytoskeletal systems into new morphological contexts (62). A third possibility is convergence, in which similar cellular constraints repeatedly favor comparable cytoskeletal adaptations despite differences in the underlying molecular repertoires (63).

The observations presented here do not distinguish among these alternatives, and all three processes may operate simultaneously. Regardless of their relative contributions, however, they operate within a broader landscape of cellular scaling constraints that favors a limited set of successful architectural solutions. Evolutionary innovation in cellular architecture is therefore shaped by historical contingency and molecular diversification, and by physical and architectural constraints that limit the range of successful solutions.

Despite these shared mechanisms, *Filoreta* and neurons differ in several fundamental respects. Neuronal arbors are typically organized as polarized branched structures centered on a soma with a single nucleus, whereas *Filoreta* develops a reticulated syncytial network through repeated anastomosis. Neuron function largely relies on directional cytoskeletal organization, while *Filoreta* distributes its nuclei and other organelles through actively coordinated transport throughout the network. These differences demonstrate that similar arborized morphologies can emerge through distinct evolutionary adaptations to common cellular constraints.

The shared mechanisms and divergent architectures observed in *Filoreta* and other arborized systems demonstrate that arborization does not represent a single conserved morphology. Similar constraints on exploration, stabilization, and long-distance transport repeatedly favor branching architectures, while differences in topology, network organization, and regulatory systems generate distinct architectural outcomes. While actin and tubulin are deeply conserved, their associated motors, nucleators, and regulatory proteins vary extensively across lineages (64–67). MyTH4-FERM myosins involved in filopodial extension evolved convergently (6,7), whereas dynein has been entirely lost in land plants and replaced by expanded kinesin repertoires (8). Arborization therefore represents not a single evolutionary innovation, but a recurrent architectural outcome that emerges when conserved cytoskeletal systems confront the challenges of increasing cellular scale. Conserved actin and microtubule systems provide a common molecular foundation, while lineage-specific diversification of motors, nucleators, and regulatory proteins permits multiple evolutionary routes to arborized cellular organization.

### Filoreta as an emerging model for evolutionary cell biology

Experimentally tractable rhizarian systems remain rare, leaving major gaps in our understanding of how cytoskeletal systems generate morphology in this deeply divergent eukaryotic lineage. To our knowledge, this study represents the first cell biological analysis of *Filoreta* and one of the first mechanistic investigations of cytoskeletal organization in Rhizaria since the foundational work on *Reticulomyxa*. As a cultivable arborized syncytium, *Filoreta ramosa* provides an emerging model for investigating cytoskeletal self-organization, intracellular transport, and the development of complex cellular architectures outside of traditional Opisthokont systems.

Several features make *Filoreta* particularly valuable as a comparative model. Its extensive arborized network develops through readily observable transitions between actin-rich exploratory protrusions and microtubule-stabilized branches, enabling direct analysis of branch initiation, stabilization, remodeling, and intracellular transport. The syncytium supports rapid long-range trafficking, distributed microtubule nucleation, and continuous network reorganization, allowing these processes to be studied within a single multinucleate cell rather than through multicellular developmental programs. The decentralized organization of the network further expands the range of experimentally accessible questions. Branch nodes function as sites of microtubule nucleation and transport coordination, while repeated anastomosis generates a reticulated topology that differs fundamentally from the hierarchical organization of neuronal arbors. Together, the combination of branching, transport, distributed nucleation, and network-level organization provides a unique opportunity to investigate how conserved cytoskeletal systems generate and maintain large-scale arborized cellular architectures.

More broadly, *Filoreta* expands the comparative toolkit available for evolutionary cell biology. By establishing a tractable rhizarian model system, *Filoreta* provides an opportunity to test how conserved cytoskeletal machinery generates evolutionary novelty while remaining constrained by the demands of cellular scale. Comparisons among Rhizaria, Amoebozoa, fungi, plants, and animals will help reveal the extent to which complex cellular architectures arise through historical contingency, evolutionary innovation, or recurring architectural principles that transcend phylogenetic boundaries.

## ACKNOWLEDGMENTS

This work was supported by the National Aeronautics and Space Administration (NASA) Astrobiology Institute through grant 12-NAI-0023 (PI: Nigel Goldenfeld, University of Illinois Urbana-Champaign) and by the National Institutes of Health, National Institute of Allergy and Infectious Diseases (NIAID), grant R01AI077571 (PI: Scott Dawson).

Additional support was provided through Whitman Center Research Awards (2016 and 2017) and a Whitman Center Nikon Fellowship (2018) to SCD from the Marine Biological Laboratory, Woods Hole, Massachusetts. We thank the faculty, staff, and students of the Marine Biological Laboratory and the Microbial Diversity Advanced Training Course for providing research facilities, training opportunities, imaging resources, technical expertise, and an intellectually stimulating scientific environment that contributed to this work. The authors also thank Sean Collins (University of California, Davis), Bo Liu (University of California, Davis), and Kassie Ori-McKenney (University of California, Davis) for valuable feedback and insightful comments on the manuscript.

## METHODS

### *Filoreta* culture conditions

*Filoreta ramosa* (strain SW4B) was isolated in September 2014 from brackish sediment samples collected in July 2014 at the Little Sippewissett Marsh in Massachusetts (41°34’33.6“N 70°38’22.7”W). The isolate was obtained by a dilution plaque plate method as follows: the environmental sample was serially diluted to 10^-3^ and suspended in a nascent bacterial culture from the same site (*Maribacter* sp.) at a density of 0.5 by McFarland Standard. The dilution was spread onto 1.5% agar plates containing Seawater Base (SWB) (423mM NaCl, 9mM KCl, 9.27mM CaCl_2_ dihydrate, 22.9mM MgCl_2_ hexahydrate, 25.5mM MgSO_4_ heptahydrate, 2.14 mM NaHCO_3_), supplemented with 0.01% Bacto yeast extract and 0.01% Bacto tryptone. The plates were incubated at 22°C and monitored for plaques. Plaques were diluted and plated as before for secondary isolation to ensure a monoeukaryotic isolate was obtained.

The strain was maintained as dormant cysts at room temperature in sterile culture flasks (Falcon™ 353135) containing filter-sterilized SWB, buffered with 5 mM MOPS. Cultures were re-grown from cysts at 22°C in 5mM MOPS-buffered SWB, enriched to a 0.01% final concentration of Yeast Extract and Tryptone. Active syncytia were maintained by refreshing enriched culture medium every 24-48 hours and were observed to fully encyst 72-96 hours after the removal of bacterial suspension and refilled with plain seawater medium. Individual amoebae were grown in shallow dishes containing 0.05% YET over 48 hours or harvested from plates when grown on a lawn of *Maribacter* sp. “prey” bacterial food.

### Generation of custom antibodies

Polyclonal rabbit antibodies were generated using Genscript’s PolyExpress Antibody services with synthetic peptide synthesis. Specific peptides derived from *Filoreta* EB1 and GCP3 homologs were selected for their location within the proteins, affinities and their predicted antigenic regions. Regions within GCP3 N-terminal end included peptides CAQSSEGSEAPPATK, TKQTSPQKPESERVC and LKRMLRNDRTKLVEC. For the EB1 homolog, we used peptide sequences CSRSSGASRKPAGTR, KSGGKPKKARGVTRC and CTRSSTSTPSRSTSG.

### Fixation and cytoskeletal immunostaining

Syncytia grown to high confluence in culture flasks were gently removed using cell scrapers (GeneMate T-2443-1). Cell suspensions were added to glass coverslips pre-coated in 0.1% cold-water fish skin gelatin (Sigma G7765), placed into separate wells of a rectangular 8-well dish (Thermo 267062). Syncytial fragments were allowed to adhere for ranges of time varying from 30 minutes to 12 hours, depending on the stage of growth to be observed (30 minutes for new fragment fusions, 1-6 hours for syncytial outgrowth, 8-12 hours for syncytial pruning, hub formation, encystation). Prior to fixation, the coverslips were submerged for five minutes in filter sterilized calcium-free artificial seawater to remove excess calcium from the medium (NaCl (469 mM), KCl (10 mM), MgCl_2_ (36 mM), MgSO_4_ (17 mM), HEPES-NaOH (10 mM, pH 8.2), EGTA (10 mM)). Three protocols were developed and used for fixation of the marine amoeba included three fixative methods: paraformaldehyde, glutaraldehyde, or glyoxal.

Paraformaldehyde (4%) fixative (20ml calcium-free seawater containing 0.4 M sucrose, 8ml cytoskeletal stabilizing buffer (CSB) (recipe below), 4ml 32% paraformaldehyde) was added for 10 minutes. CSB was prepared as a 10x stock and contained 50 mM KCl, 1.37 M NaCl, 40 mM NaHCO3, 110 mM Na2HPO4, 20 mM MgCl2, 50mM PIPES, 20 mM EGTA. Glutaraldehyde fixative (0.5% glutaraldehyde, 5% sucrose, in calcium-free seawater buffered with 1X CSB) was added for one minute. Glyoxal fixative (1.5ml of 40% glyoxal (Sigma 128465), 2ml ethanol, 1ml CSB, 7ml calcium-free seawater) was added for 30 seconds. Each fixation was quenched by three successive two-minute washes in PEM (100 mM PIPES, 1 mM EGTA, 0.1 mM MgSO4) (68), followed by a 10-minute permeabilization in 0.1% Triton-X 100, blocked in PEMBALG (PEM containing 1% bovine serum albumin, 0.1% sodium azide, 100 mM lysine, and 0.5% cold-water fish skin gelatin (Sigma G7765)) (68). In glutaraldehyde fixation, 0.2 M Glycine was added for 30 minutes to quench unreacted aldehydes to reduce autofluorescence.

Primary antibodies were incubated on coverslips at optimized concentrations (Mu-ɑ-TAT1= 1:500), Rb-ɑ-GCP3=1:1000 (Custom antibodies GenScript PolyExpress Gold – *Filoreta* GCP3 peptides), Rb-ɑ-Ɣ-Tubulin=1:1000 (Abcam ab205475) for one hour at room temperature, then washed three times in PEMBALG prior to incubation with the secondary antibody Alexa fluor 488, 594, 647 conjugates(1:1000) for the same duration at room temperature in a dark cabinet. Actin networks were labeled using a 1:1000 concentration of Phalloidin-AlexaFluor 488 conjugate in PEM for 10 minutes (PFA or Glutaraldehyde fixation only). Nuclei were labeled by counterstaining in a 1:1000 dilution of DAPI in PEM for 10 minutes in a dark cabinet. Slides were mounted with (SlowFade™ Diamond Antifade Mountant S36963). Multichannel acquisition of stained syncytia was performed with widefield microscopy on a Leica DMI6000B with filter cube set N3, A4, L5, Cy5 and captured using a Prime 95B sCMOS camera. Images were analyzed in FIJI (ImageJ) (69), and maximum intensity projections (MIPs) were generated for structural analysis.

### Live differential interference contrast (DIC) imaging

Syncytia grown to high confluence in surface treated culture flasks were gently rinsed in sterile medium, scraped, and suspensions were added to 30mm Mattek dishes with glass coverslips coated with 0.1% gelatin. Syncytial fragments were allowed to adhere for ranges of time varying from 30 minutes to 12 hours, depending on the stage of growth to be observed (30 minutes for new fragment fusions, 1-6 hours for syncytial outgrowth, 8-12 hours for syncytial pruning, hub formation, encystation). Live imaging used differential interference contrast (DIC) with a Leica DMI6000B with shutter set to “open” on low light levels (10%) for growth and shutter set to “auto” for pruning. Timelapse movies were captured at one second intervals.

### Sholl analysis of arborization

Discrete time points were isolated from imaging data and processed prior to quantification of arborization. DIC images were background cleaned in FIJI, where a duplicate image is Gaussian blurred, then divided against the original image to remove artifact background shading. Ideal thresholding levels varied by gray values across different image series, but each time point in a given series used identical thresholding parameters. Excess pixel noise was removed using “Analyze Particles” to generate masks for objects smaller than 10 pixels. The binary images corresponding to time points in a series were synchronized and identically annotated with a line drawn from a central point (identified as a syncytial hub or major branch node) towards the periphery of the image. Identical parameters were used in FIJI’s Sholl Analysis software across all treatments. Sholl Intersection Profiles (SIPs) generated by the software were exported and used in Prism GraphPad to extrapolate further values in “Growth rate” and “Area under curve”. The intersection number within serial spheres of 1 μm radii was plotted as a function of radial distance from the center. Plots generated from images within the same time series were plotted together, and SIP peaks were used to quantify the change in network diameter over time.

### Branch growth rate measurements

Sholl profiles were exported from FIJI for use in GraphPad Prism Software. Profiles were plotted along an X-Y graph, and the Y-value (number of intersecting branchlets) of the half-maximal peak for each Sholl profile in a series (0 min, 10 min, 20 min) was taken with its corresponding X-value (distance from center). The differences in these distances were taken and divided by minutes in the time series to generate a table of micrometer distance over one minute. Additional calculations included averaging growth measurements taken at one-minute intervals and averaged to ensure rates were not artificially elevated by using larger time intervals.

### Area under curve measurements

GraphPad Prism Software analyses were used to quantify total surface area coverage by the syncytial network by generating “Area Under Curve” values for XY corresponding datasets. The area under curve data were generated using only raw values and not the “best fit” polynomial regressions produced in Sholl profiles. These AUCs were then compared across timepoints as a ratio to the original T=0 timepoint for each replicate, divided by time elapsed (for “per minute” values). Values increasing in AUC are reported as increased overall surface area (branch intersection numbers are maintained and/or increasing in number as the syncytial growth rate increases). Values decreasing in AUC are reported as decreased surface area (pruning/simplification, loss of branches, destabilization, or pseudopodial retraction).

### Timelapse Sholl analysis of arborization

While each syncytial growth experiment was treated identically prior to imaging, there was no normalization of bacterial food presence in cultures other than the specific concentration of nutrient source provided. Since day-to-day imaging incorporated these potential sources of variation, we color coded data points by experimental day to indicate variation. Live imaging of syncytia was performed at one second intervals (frame rate), and analyses used five- and ten-minute intervals for comparison of control and drug-treated growth rates and topology maintenance. Images were thresholded with identical parameters within each replicate. Sholl profiles generated in FIJI were compared for growth rate (distance to most peripheral pseudopod radial intersection) and topology (area under curve, “total peak area” of datapoints graphed in GraphPad Prism Software).

### Fluorescence Sholl analysis of arborization

Images were prepared from raw data by thresholding maximum intensity projections (MIPs) using Fiji (normalized to each color for comparative 2-channel fluorescence Sholl). Center points were selected as the main nodes for multiple branches in broad scale syncytia, and main branch nodes for peripheral branchlet arbors, respectively. The comparison of actin and microtubule Sholl profiles used the ratio of total peak area for each channel’s profile, in which branch intersections were normalized to each acquisition by using identical thresholding and Sholl profile parameters. The total area of microtubule SIPs were divided by the total area of actin SIPs to find the percentage of microtubule proliferation within the actin network. Identical parameters were used across drug treatments.

### Quantification of organelle transport

Syncytia were stained in 96-well glass-bottomed plates, and in Mattek dishes pre-coated with 0.1% gelatin. Nuclei, mitochondria, and lysosomes were stained for 10 minutes at room temperature with Hoechst 33342, Syto9, and LysoTracker, respectively. Nuclei were imaged at 1 second intervals (50 ms exposure, 10% excitation power) under widefield fluorescence, while mitochondria and lysosomes were able to be imaged at 100 millisecond intervals.

Imaging experiments incorporated the control for each field of view imaged by capturing the first 60 seconds without drug, then added the drug or buffer at equal volume for each replicate. Ciliobrevin D (Fisher cat# 25040110MG), Latrunculin A (Fisher cat# L12370) and Nocodazole (Fisher cat# 50-165-6973) were used at the final concentrations indicated in figure legends. Syncytia that showed signs of phototoxicity within the 5 minutes of fluorescent light exposure (blebbing, beading, shriveling) were excluded from data collection to exclude the stress response altering biologically relevant values.

Organelle trajectories were quantified using kymographs generated in FIJI plugin “KymoResliceWide” from manually traced branches and verified using particle tracking software in the FIJI plugin “Trackmate” with automated settings. Rates of transport were calculated from kymographs using the line tool, for processive movements of organelles moving 10 µm or more. Organelles moving fewer than 10 µm were excluded from the dataset. Rate measurements were then added to superplots in Prism GraphPad and colorized per replicate (70). Drug-treated image series of organelle traffic kymographs were generated using identical parameters to their controls and compared between series’ concentrations for each data set within GraphPad using ordinary one-way ANOVA and multiple comparison analyses compared against the control.

### Gamma-tubulin, and GCP3 immunostaining and analysis

Immunostaining and localization of microtubule interacting proteins were visualized using custom polyclonal antibodies against *Filoreta* GCP3 peptide #3 (LKRMLRNDRTKLVEC). Syncytia were grown and fixed on 0.1% gelatin-coated coverslips, then fixed using glyoxal and paraformaldehyde fixation and immunostaining methods outlined above. Tubulin was stained with mouse-anti-alpha-tubulin antibody at a 1:500 ratio in PEMBALG (‘TAT-1’, an inherited gift from Keith Gull (71). The Alexa Fluor 488 Anti-gamma-tubulin antibody (Abcam cat# ab205475) and GCP3 antibodies were used at a 1:1000 ratio in PEMBALG for 1 hour at 22°C or 4°C overnight. Secondary antibodies against the rabbit-GCP3 and mouse-TAT-1 were added at a 1:1000 ratio, using goat-anti-rabbit AlexaFluor647 conjugate and goat-anti-mouse AlexaFluor 594 conjugates for 1 hour at 22°C. Syncytia were counterstained with DAPI and mounted with SlowFade™ Diamond Antifade Mountant S36963. Images were taken on a Leica DMI6000B with Prime95B sCMOS camera.

The resultant co-stained images were analyzed using FIJI using measurement features over 15-pixel wide segmented lines. The average grey values were calculated over branches and nodes and normalized to length and width of areas covered by branches and nodes. Each branch thickness was measured using the “plot profile” function and used to categorize measurements for “minor” and “major branches” compared to “nodes” which included three or more branch intersections. The grey values/area were plotted for each image using Prism GraphPad and colored per replicate.

